# Measuring Robustness of Brain Networks in Autism Spectrum Disorder with Ricci Curvature

**DOI:** 10.1101/722025

**Authors:** Anish K. Simhal, Kimberly L.H. Carpenter, Saad Nadeem, Joanne Kurtzberg, Allen Song, Allen Tannenbaum, Guillermo Sapiro, Geraldine Dawson

## Abstract

Ricci curvature is a method for measuring the robustness of networks. In this work, we use Ricci curvature to measure robustness of brain networks affected by autism spectrum disorder (ASD). Subjects with ASD are given a stem cell infusion and are imaged with diffusion MRI before and after the infusion. By using Ricci curvature to measure changes in robustness, we quantify both local and global changes in the brain networks correlated with the infusion. Our results find changes in regions associated with ASD that were not detected via traditional brain network analysis.

---

The prevalence of neurodevelopmental disorder diagnosis, such as autism spectrum disorder (ASD), has been growing over the past few decades. According to one recent study, almost 17% of children in the United States have been diagnosed with a neurodevelopmental disorder and approximately 2.5% are diagnosed with ASD [1]. ASD is clinically characterized by restricted interests and repetitive behaviors as well as social communication deficits [2]. Individuals with ASD have atypical white matter developmental patterns compared those typically developing subjects and this difference is linked to the severity of ASD symptoms [3, 4]. White matter development can be altered by neuroinflammation, which in turn, is associated with abnormalities in cerebrospinal fluid circulation [5]. Autologous cord blood infusions, a potential therapy, are theorized to reduce neuroinflammation [6, 7] and promote white matter development, thus triggering a reconfiguration of connectivity patterns in the brain [8]. In a previous study, improvements in social functioning and communication abilities following treatment were correlated with an increase in connectivity of the white matter networks underlying the social and communicative functions [8, 9]. These changes in both white matter volume and connectivity are usually measured via diffusion tensor imaging (DTI), a form of magnetic resonance imaging (MRI) which measures the diffusion of water molecules throughout the brain, a correlate for brain connectivity. While exploring these changes is a useful measure, it does not account for all the connections into the brain region or pairs of brain regions or examine the robustness of connections in the brain network. To take advantage of the network imaged by DTI, we need a measure that reflects a specific brain region’s relationship with every other region in the brain and that quantifies the robustness of such broad network connections. This is addressed in this work.

One solution to measure the robustness of brain regions is Ricci curvature. Broadly speaking, curvature is a “measure by which a geometrical object deviates from being flat” [10]. Although there are multiple notions of graph curvature [11], this work focuses on the Ricci curvature as formulated by Ollivier [14], because of its positive correlation with the robustness of a network and because of its natural physical interpretation and computational efficiency. The link between curvature and robustness is as follow: curvature correlates positively with entropy; entropy correlates positively with robustness [10]. Robustness measures the extent to which a network can withstand perturbations. For brain networks, robustness measures the extent a region in a brain or a connection between two regions in the brain can be affected or withstand damage by a disease or a treatment.

The formal connection of curvature to robustness arises from several sources, including systems and control theory. Feedback tends to make a given system less sensitive, i.e., more robust, to parameter variations and external disturbances [12]. For a weighted graph derived from DTI, feedback is represented by the number of invariant triangles at a given node. Therefore, the greater the number of triangles [13], the higher the curvature value. The curvature between two brain regions is computed by using a distance derived from the theory of optimal transport, and gives a novel measure of connectivity and feedback stability based on both local and global network geometry [14]. Thus, the curvature between two brain regions considers the strength of connection between those two brain regions and also the rest of the brain. This measurement takes into account the context of a brain region pair and serves as a useful lens through which to analyze the robustness of the brain.

To measure the efficacy of cord blood infusions for improving social functions of children with ASD for a pilot clinical trial, nineteen participants were imaged via DTI and participated in a series of behavioral exams before and after the treatment. A full characterization of the sample is provided in [8]. The brain regions for each subject were delineated and defined as nodes of a network, while edges described structural connectivity between them. The DTI parcellation was done using the UNC Pediatric Brain Atlas. Ricci curvature was computed for the edges and a version of scalar curvature at the nodes by taking the weighted average of the Ricci curvature over all the neighboring edges. This is described in detail in the methods section. For these subjects, behavioral tests were administered before and after the treatment as well. When analyzing the data, we looked at the change in behavioral scores and change in curvature. These changes were compared via Pearson correlation. The results presented are the changes in curvature between two nodes which correlate significantly (p < 0.05) with the change in behavior with two or more clinical tests. Potentially due to the relatively small data sample, none of the scalar (node) or edge (connection between ROIs) curvature correlations survived a false discovery rate (Benjamini-Hochberg correction) with a p-value of 0.05.

The results highlight regions which have been previously indicated in ASD, but were not evident when constrained to the differences in white matter connectivity between pairs of individual brain regions. Furthermore, these differences were not found when using traditional brain network analysis techniques, as listed in [15]. First, using Ricci curvature, we see a relationship between clinical improvement and altered robustness in white matter pathways that are implicated in the social and communication abilities that improved following treatment. For example, we demonstrate a relationship between clinical improvement across all three measures and increased curvature (robustness) in a white matter pathway connecting the right dorsolateral prefrontal cortex (dlPFC) to the right insula, as shown in Table 1 and Figure 1. Both of these regions have been implicated in autism [16, 17], with the insula in particular serving as a key structural and functional brain hub across health and disease [18]. Resting state MRI (rsMRI) studies have suggested a role of the insula in one of the three canonical rsMRI networks, the *salience network*, which plays a critical role in detecting salient information from the sensory environment and engaging other functional networks, including the central-executive network of which the dlPFC is part [19]. The central-executive network is then responsible for integrating multiple cognitive processes, including working memory and attentional control, in the support of goal directed behaviors. Further, rsMRI studies of these networks have demonstrated aberrant connectivity in these canonical networks correlates with social and communication abilities in children with ASD [20]. In addition to identifying novel white matter pathways, the curvature analysis also identified regions for which the robustness of their connections with the rest of brain increased in relation to clinical outcomes: the left fusiform gyrus and the right pars orbitalis, as shown in Table 2. Both the fusiform gyrus and the pars orbitalis, which lies within the inferior frontal gyrus (IFG), are key components of the social brain network [21]. Previous research has linked the structure and function of both the fusiform gyrus and IFG to social cognition in autism [22, 23]. Importantly, none of these white matter networks were identified using conventional DTI analyses [9].

**Table 1.** Table of edge curvature results.. Table showing the node pairs where the change in behavioral scores and change in curvature correlate with a *p* < 0.05 for two or more behavioral exams. For each edge pair listed, the associated change in curvature is listed as mean ± standard deviation. Change in behavior is measured the behavior measured at the end of the study minus the behavior measured at the beginning of the study. Change in curvature is measured as ratio of the curvature of a node at the end of the study over the curvature of the node at the beginning of the study. VABS: Vineland Adaptive Behavior Scales-II, EOW: Expressive One-Word picture vocabulary test, CGI: Clinical Global Impression Scales.

| Edge Pairs | $\Delta VABS$ | $\Delta EOW$ | $\Delta CGI$ | $\Delta Curvature$ |
| --- | --- | --- | --- | --- |
| R. Frontal Pole - R. Inferior Temporal | $3.37 \pm 6.71$ | $4.95 \pm 6.34$ | $2.79 \pm 1.82$ | $1.25 \pm 1.49$ |
| R. Rostral Middle Frontal - R. Insula | $3.37 \pm 6.71$ | $4.95 \pm 6.34$ | $2.79 \pm 1.82$ | $1.14 \pm 0.86$ |
| R. Bankssts - R. Accumbens Area | $3.37 \pm 6.71$ | $4.95 \pm 6.34$ | $2.79 \pm 1.82$ | $1.104 \pm 0.46$ |
| R. Putamen - R. Pallidum | $3.37 \pm 6.71$ | $4.95 \pm 6.34$ | $2.79 \pm 1.82$ | $0.98 \pm 0.11$ |
| L. Lateral Orbitofrontal Gyrus - L. Inferior Temporal Gyrus | $3.37 \pm 6.71$ | $4.95 \pm 6.34$ | $2.79 \pm 1.82$ | $1.83 \pm 6.85$ |
| L. Rostral Anterior Cingulate - L. Hippocampus | $3.37 \pm 6.71$ | $4.95 \pm 6.34$ | $2.79 \pm 1.82$ | $0.91 \pm 1.0005$ |

**Table 2.** Table of scalar curvature results. Table showing the nodes where the change in behavioral scores and change in curvature correlate with a *p* < 0.05 for two or more behavioral exams. For each node listed, the associated change in curvature is listed as mean ± standard deviation. Change in behavior is measured the behavior measured at the end of the study minus the behavior measured at the beginning of the study. Change in curvature is measured as ratio of the curvature of a node at the end of the study over the curvature of the node at the beginning of the study. VABS: Vineland Adaptive Behavior Scales-II, EOW: Expressive One-Word picture vocabulary test, CGI: Clinical Global Impression Scales.

| Node | ΔVABS | ΔEOW | ΔCGI | ΔCurvature |
| --- | --- | --- | --- | --- |
| Right Pars Orbitalis | $3.37 \pm 6.71$ | $4.95 \pm 6.34$ | $2.79 \pm 1.82$ | $1.0042 \pm 0.0595$ |
| Left Pericalcarine | $3.37 \pm 6.71$ | $4.95 \pm 6.34$ | $2.79 \pm 1.82$ | $1.0099 \pm 0.083$ |
| Left Fusiform | $3.37 \pm 6.71$ | $4.95 \pm 6.34$ | $2.79 \pm 1.82$ | $1.022 \pm 0.121$ |
| Left Transverse Temporal Gyrus | $3.37 \pm 6.71$ | $4.95 \pm 6.34$ | $2.79 \pm 1.82$ | $1.015 \pm 0.0722$ |

**Figure 1.**
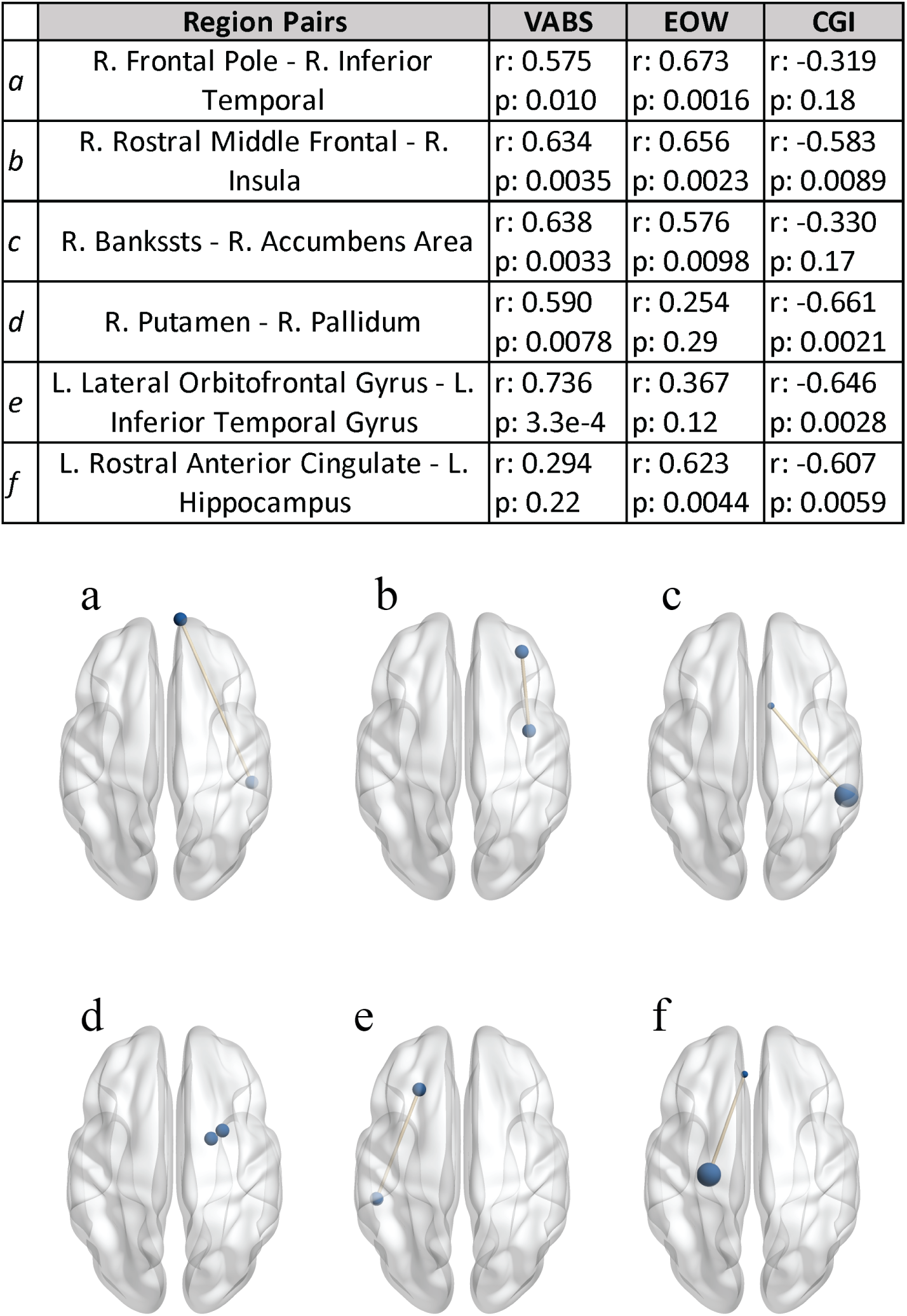
Overview of the study and results. Top) Table showing the brain region pairs and the Pearson correlation coefficient between the change in curvature and change in behavioral scores. The associated p-value is reported below the correlation coefficient. Bottom) Axial projection of pairs where the change in curvature correlated significantly with behavioral exams. Cross hemisphere brain connections were removed for this analysis. The brain graphics were visualized with the BrainNet Viewer (http://www.nitrc.org/projects/bnv/) [38]. VABS: Vineland Adaptive Behavior Scales-II, EOW: Expressive One-Word picture vocabulary test, CGI: Clinical Global Impression Scales.

These results must be considered in light of some limitations. Because this was an open label trial, it is not possible to determine whether the clinical and curvature changes were a result of normal trajectories of improvement and development or whether they were a consequence of the treatment itself. However, the current data provides targets for exploring brain-related changes in future placebo-controlled double-blind trials, which are currently taking place. Due to the number of DTI directions that were captured and the resolution of the data, cross hemisphere brain connections were removed. Future studies using higher-dimensional data are warranted.

Despite significant progress in understanding the underlying neurobiology of ASD, there are still few reliable and objective measures of change in social and communication function ASD and their relationship with underlying brain structures. We show that curvature identifies changes in robustness in brain regions that may be attributed to the changes in brain connectivity caused by the cord blood infusion. Thus, this study lays the foundation for a new approach to assess both the robustness of a specific brain region and brain region pairs.

## Methods

### Study Design and Sample

The current study is a secondary data analysis of DTI data collected as part of a phase 1 open-label trial of a single intravenous infusion of autologous umbilical cord blood in 25 children with ASD who were between 24-72 months of age at baseline. The methods of this trial and the accompanying DTI analyses have been described in detail elsewhere [8, 9, 24, 25]. Children with a confirmed diagnosis of ASD and a banked autologous umbilical cord blood unit of adequate size and quality participated in the trial. Nineteen participants provided high quality, artifact-free data for the DTI at both baseline and 6-month visits (17 males and 2 females). All caregivers/legal guardians of participants gave written, informed consent, and the study protocol was approved by the Duke University Health System Institutional Review Board. Methods were carried out in accordance with institutional, state, and federal guidelines and regulation. All methods and the trial were approved by the Duke Hospital Institutional Review Board and conducted under IND #15949. The ClinicalTrials.gov Identifier is NCT02176317.

### Clinical Measures

Social abilities were measured with the Vineland Adaptive Behavior Scales-II Socialization Subscale (VABS-SS) [26]. The change in the VABS-SS (6 month-baseline) was used to measure change in social behavior. The Expressive One Word Picture Vocabulary Test 4 (EOW) is a clinician-administered assessment which measures an individual’s ability to match a spoken word with an image of an object, action, or concept [27]. The change in the raw score (6 month-baseline) was used to measure change in expressive language. Finally, overall clinical severity and improvement was measured with the Clinical Global Impression Severity (CGI-S) and Improvement (CGI-I) scales.

### Magnetic Resonance Imaging Acquisition and Analysis

MRI scanning was conducted on a 3.0 T GE MR750 whole-body 60cm bore MRI scanner (GE Healthcare, Waukesha, WI). Participants were sedated to reduce motion artifacts in the MRI. Diffusion weighted images were acquired using a 25-direction gradient encoding scheme at *b* = 1000*s/mm*^2^ with three non-diffusion-weighted images, an average (std) echo time (TE) of 85*m*s (2*ms*), and a repetition time (TR) of 12, 000*ms*. An isotropic resolution of 2*mm*^3^ was achieved using a 96 × 96 acquisition matrix in a field of view (FOV) of 192 × 192*mm*^2^ at a 2*mm* slice thickness. T1-weighted images were obtained with an inversion-prepared 3D fast spoiled-gradient-recalled (FSPGR) pulse sequence with a TE of 2.7*ms*, an inversion time (TI) of 450*ms*, a TR of 7.2*ms*, and a flip angle of 12°, at a 1*mm*^3^ isotropic resolution.

### Connectome Analysis Pipeline

The full connectome analysis pipeline is described in detail elsewhere [9]. Briefly, each participant’s T1 image and the first non-diffusion weighted image (b0) of the DTI acquisition were skull-stripped using the FSL brain extraction tool [28, 29]. The T1 image was registered to the b0 image with an affine registration created using FSL FLIRT [30, 31]. Region of interest (ROI) parcellation was performed by warping the dilated UNC Pediatric Brain atlas (available publicly at http://www.nitrc.org/projects/unc_brain_atlas/) into each participant’s T1 in diffusion image space via the Advanced Normalization Tools (ANTs) toolkit [32, 33]. A total of 83 regions were defined for each participant, 41 gray matter regions in each hemisphere, and a single region encompassing the brainstem. FMRIB’s Automated Segmentation Tool (FAST) was used to calculate whole brain white matter volume for each participant at both baseline and 6 month visits [34]. Following this, a standardized pipeline for deterministic tractography based on the Connectome Mapper (CMP) was used to analyze participant data at both baseline and 6 month visits (http://www.cmtk.org) [35, 36]. The parcellated gray matter ROIs included in this analysis are defined as nodes. Edges are defined as the volume of voxels containing valid streamlines that originate and terminate within a pair of nodes. For each participant, edge volumes were calculated and normalized by whole-brain white matter volume at both baseline and 6-month visits.

### Curvature Analysis

In this section, we outline how we compute the Ricci curvature on discrete metric measure spaces including weighted graphs. The motivation of the Olliver-Ricci [14] definition of curvature on a weighted graph is based on the following characterization of Ricci curvature from Riemannian geometry [37]. For *X* a Riemannian manifold, consider two very close points *x, y* ∈ *X* and two corresponding small geodesic balls. Positive curvature is reflected in the fact that the distance between two balls is less than the distance between their centers. Similar considerations apply to negative and zero curvature.

For this work, the brain network is represented as an undirected and positively weighted graph, *G* = (*V, E*), where *V* is the set of *n* vertices (nodes) in the network and *E* is the set of all edges (links) connecting them with weights {*w*}. Consider the graph metric *d* : *V* × *V* → ℝ^+^ on the set of vertices *V* where *d*(*x, y*) is the number of edges in the shortest path connecting *x* and *y*. (*d* may be any intrinsic metric defined on *V*.) We let denote *w*_*xy*_ > 0 denoting the weight of the edge between node *x* and *y*. (If there is no edge, then *w*_*xy*_ = 0.) For any two distinct points *x, y* ∈ *V*, the Ollivier-Ricci (OR) curvature is defined as

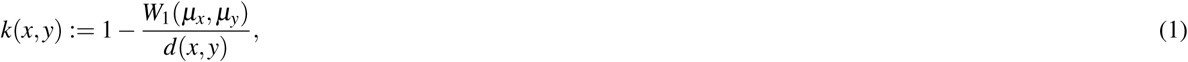

where *W*_1_ denotes the Earth Mover’s Distance (Wasserstein 1-metric). We define the weighted degree at node *x, d*_*x*_ as

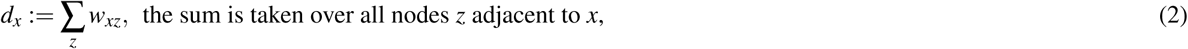

and we define the probability measure at *x, µ*_*x*_ as

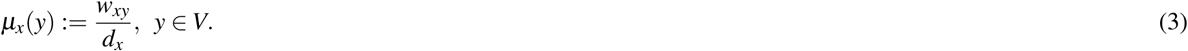

The *scalar curvature* at a given node *x* (the contraction of Ricci curvature) is defined as

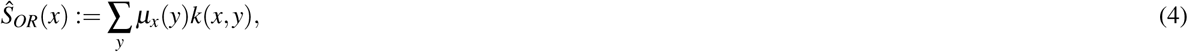

where the sum is taken over all neighbors of *x*. Given the discussed, positive correlation of robustness and curvature, in our work, we propose to use curvature as a proxy for robustness. Various advantages of using Ricci curvature in this framework are described in more detail in [10]. The code used for this analysis is shared at https://github.com/aksimhal/Curvature-ASD-Analysis.

## Acknowledgements

This work would not have been possible without the participation of the participants and their families. We thank the Marcus Foundation (NCT02176317), PerkinElmer, Inc., and the Stylli Translational Neuroscience Award for their financial support for this study, the children who participated and their families, and the following staff members: Jennifer Baker, Colleen McLaughlin, Rebecca Durham, Barbara Waters-Pick, Todd Calnan, Crystal Chiang, Kendyl Cole, Michelle Perry, Mallory Harris, Jennifer Newman, Katherine S. Davlantis, Elizabeth Paisley, Charlotte Stoute, and Elizabeth Sturdivant. This work was supported by the following: National Science Foundation (NSF), United States Office of Naval Research (ONR), United States Army Research Office (ARO), United States Air Force Office of Research (AFOSR), National Institute of Aging (NIA), Breast Cancer Research Foundation (BCRF), and the National Geospatial-Intelligence Agency (NGA). Gifts to G.S. from Google, Microsoft, and Amazon are also acknowledged.

## Author Contributions

AKS, KC, SN, AT, GS, and GD designed the study and wrote the paper. AS is in charge of the MRI data collection. JK leads the blood cord effort. JK and GD designed the blood cord intervention in ASD. AKS and SN implemented the algorithms. AKS, KC, and GD analyzed the results.

## Author Declarations

G.D. is on the Scientific Advisory Board and receives research funding from Janssen Research and Development, LLC, is a consultant for Roche Pharmaceuticals, LabCorps, Inc., Apple, Gerson Lehrman Group, Guidepoint, and Akili, Inc., received research funding from PerkinElmer and Janssen, and receives royalties from Guilford Press, Springer and University of Oxford Press. J.K. is Director of the Carolinas Cord Blood Bank and Medical Director of Cord:Use Cord Blood Bank. G.S. consults for Apple Inc., and Volvo Cars, and is on the Board of Directors of SIS.

## Supplemental Curvature Information

### Introduction

Geometric techniques can be used to study the properties of complex networks, including as a method to measure a network’s robustness to certain perturbations [1, 2, 3, 4, 5]. Curvature, for this task, is based on the theory of optimal mass transport [6, 7]. First, note that the space of probability densities on a given Riemannian manifold also inherits a natural Riemannian structure [8, 9, 10] for which changes in entropy and Ricci curvature are positively correlated [11, 12]. In conjunction with the Fluctuation Theorem [13], we can conclude that an increase in Ricci curvature is positively correlated with an increase in network robustness, herein expressed as ΔRic × Δ*R* ≥ 0. This is the key observation that we will exploit below. This observation leads to a notion of curvature on the space of probability distributions on rather general metric measure spaces.

There have been several notions of graph curvature proposed, including those based on the Bochner-Weitzenböck decomposition of the Laplace operator [14] (Forman-Ricci curvature), and those based on Bakry-Émery theory [5, 15]. In this work, we use the formulation of Ollivier [16] because of its direct link to the geometry of optimal mass transport, and therefore its link to entropy and network robustness. There are several other properties of the Ollivier-Ricci curvature that make it attractive for studying the robustness of networks. There is a direct correlation of changes in the rate function from large deviations theory and Ollivier-Ricci curvature [1, 16]. Second, the Ollivier-Ricci curvature is a natural measure of feedback connectivity. This is because the number of triangles in a network (redundant pathways) can be characterized by an explicit lower bound based on Ollivier-Ricci curvature [17]. Similar to control theory, redundancy in feedback is the main characteristic of a robust system architecture. Finally, positive Olliver-Ricci curvature governs the rate of convergence to an invariant distribution on a weighted graph which is modeled as a Markov chain. For these reasons, we chose to use the Ollivier-Ricci curvature for our analysis.

### Ricci curvature and entropy

We briefly sketch the connection of curvature and entropy and curvature on a Riemannian manifold. Accordingly, let *X* denote a complete connected Riemannian manifold equipped with metric *d*. Ricci curvature provides a way of measuring the degree to which the geometry determined by a given Riemannian metric differs from that of ordinary Euclidean space [18]. The Ricci curvature tensor is obtained as the trace of the sectional curvature [18], and a lower bound on Ricci curvature provides an estimate of the tendency of geodesics to converge or diverge. Through the work of Lott-Sturm-Villani [11, 12], optimal transport connects the bounds on Ricci curvature with entropy, as sketched out below.

First, let *𝒫*(*X*) denote the space of probability densities with finite second moments on *X*. One can show that *𝒫*(*X*) also has a Riemannian structure [8, 10] via the Wasserstein metric *W*_2_ from optimal mass transport theory (OMT). Next, for a probability measure *µ* ∈ *𝒫*(*X*), the *Boltzmann entropy* is defined as

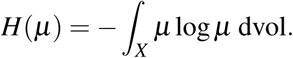

Lott, Sturm, and Villani [11, 12] found a very interesting connection between Ricci curvature and the Boltzmann entropy via OMT. Namely, the Ricci curvature Ric ≥ *k* if and only if the entropy functional is displacement *k*-concave along the 2-Wasserstein geodesics, that is, for *µ*_0_, *µ*_1_ ∈ *𝒫*(*X*) we have

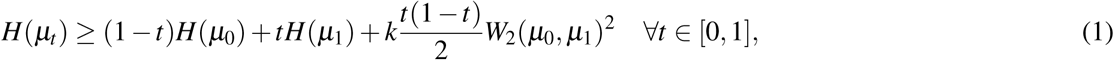

where (*µ*_*t*_)_0≤*t*≤1_ is a 2-Wasserstein geodesic between *µ*_0_ and *µ*_1_. This inequality indicates the positive correlation between changes in entropy and changes in curvature, which we denote as

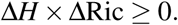

### Entropy and robustness

As previously mentioned, we are interested in the connection between robustness and the network entropy. Informally, functional robustness is defined as the ability of a system to adapt to random perturbations in its environment. This has been exploited in several works [1, 2, 3, 4] in order to employ changes in curvature as a proxy for robustness. This has been extensively explored in the aforementioned papers, and so we will only briefly describe the necessary results.

The starting point is the paper [13], in which the authors characterize robustness via the fluctuation decay rate after random perturbations. Denote by *p*_*ε*_ (*t*), the probability that the deviation of the sample mean is more than *ε* at time *t* from the original value. Following large deviations theory [19], the *fluctuation decay rate* is defined as

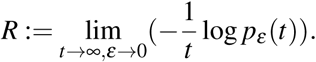

The larger the value of *R*, the faster the systems converges to a stationary state. The Fluctuation Theorem [13] states that the changes in entropy and robustness are positively correlated:

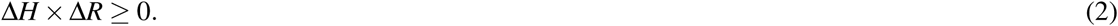

Combining (1) and (2) give a relationship which indicates the correlation of changes in curvature and robustness:

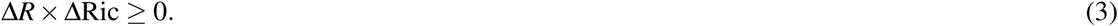

Thus changes in curvature serve as a measure of changes in system robustness. Positive changes indicate increase in robustness, while negative changes indicate the increase in system fragility.

### Wasserstein distance on discrete spaces

We will not need a general definition of the Wasserstein distance in defining the curvature, and so we will suffice with a definition that applies to a weighted connected undirected graph *G* = (*V, E*) with *n* nodes (*V*) and *m* edges (*E*). See [6, 7, 20] and the references therein for the details.

Given two probability densities *ρ*^0^, *ρ*^1^ ∈ ℝ^*n*^ on the graph, the Kantorovich formulation of the optimal transport problem [6, 7] seeks a joint distribution *ρ* ∈ ℝ^*n*×*n*^ with marginals *ρ*^0^ and *ρ*^1^ minimizing the total cost Σ *c*_*i j*_*ρ*_*i j*_:

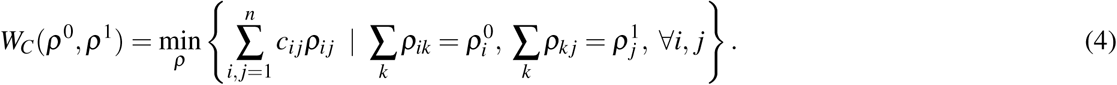

Here, *c*_*i j*_ is the cost of moving unit mass from node *i* to node *j*. If *c*_*i j*_ is defined via some intrinsic distance on *G* (e.g., the hop metric), the minimum of (4) defines a metric *W*_1_ (the Earth Mover’s Distance) on the space probability densities on *G*.

An alternative formulation may be formulated via the fluxes *u* ∈ ℝ^*m*^ on the edges. Letting *D* ∈ ℝ^*n*×*m*^ denote the oriented incidence matrix of *𝒢*, we have

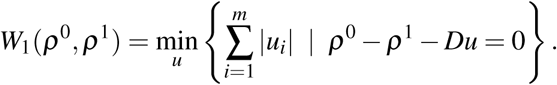

The incidence matrix *D* = [*d*_*ik*_] ∈ ℝ^*n*×*m*^ is defined by associating an orientation to each edge *e*_*k*_ = (*i, j*) = (*j, i*) of the graph: one of the nodes *i, j* is defined to be the head and the other the tail, and then we set *d*_*ik*_ = +1(−1) if *i* is the head (tail) of *e*_*k*_ and 0 otherwise. Compared to the Kantorovich formulation which has *n*^2^ variables, the above formulation has only *m* variables. It may greatly reduce the computational load when the graph *G* is sparse, i.e., *m* << *n*^2^ and is often the case in real data.

### Olliver-Ricci curvature

In this section, we define the Ricci curvature on discrete networks. Specifically, we assume that our network is presented by an undirected and positively weighted graph, *G* = (*V, E*), where *V* is the set of *n* vertices (nodes) in the network and *E* is the set of all edges connecting them, with *w*_*xy*_ > 0 denoting the weight of the edge between node *x* and *y*. (If there is no edge connecting the two nodes, then *w*_*xy*_ = 0.) We should note that in our treatment above we connected curvature and entropy via *W*_2_, and now we will be using *W*_1_. This technical point is discussed and justified in detail in [3].

Let *d*_*x*_ := Σ_*z*_ *w*_*xz*_, where the sum is taken over all nodes *z* in a neighborhood of *x*. At each node, we define the probability measure

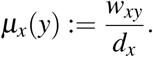

Let *d* : *V* × *V* → ℝ^+^ denote an intrinsic metric on *G*, e.g., the hop distance. Then for any two distinct points *x, y* ∈ *V*, the *Ollivier-Ricci (OR) curvature* [16] is defined as

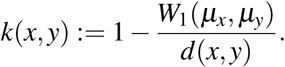

Using this edge based notion of curvature, we can define the *scalar curvature* at a given node by

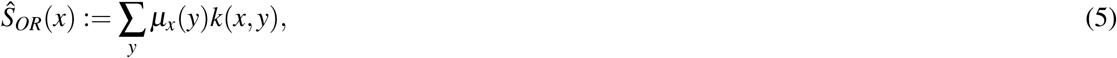

where the sum is taken over all neighbors of *x*.

Given our above arguments, the positive correlation of robustness and curvature, we use curvature as a proxy for robustness in our study of autism. Curvature can be defined either *nodally* (scalar) or relative to *edges* (Ricci). Various advantages of using Ricci curvature in this framework are described in more detail in [1].

